# Diverse biological processes coordinate the transcriptional response to nutritional changes in a *Drosophila melanogaster* multiparent population

**DOI:** 10.1101/712984

**Authors:** E. Ng’oma, P.A. Williams-Simon, A. Rahman, E.G. King

**Author notes:** **Corresponding Author:** Enoch Ng’oma, Postdoctoral Scholar, 401 Tucker Hall, University of Missouri, Columbia, MO 65211, USA.

## Abstract

**Background:** Environmental variation in the amount of resources available to populations challenge individuals to optimize the allocation of those resources to key fitness functions. This coordination of resource allocation relative to resource availability is commonly attributed to key nutrient sensing gene pathways in laboratory model organisms, chiefly the insulin/TOR signaling pathway. However, the genetic basis of diet-induced variation in gene expression is less clear.

**Results:** To describe the natural genetic variation underlying nutrient-dependent differences, we used an outbred panel derived from a multiparental population, the *Drosophila* Synthetic Population Resource. We analyzed RNA sequence data from multiple female tissue samples dissected from flies reared in three nutritional conditions: high sugar (HS), dietary restriction (DR), and control (C) diets. A large proportion of genes in the experiment (19.6% or 2,471 genes) were significantly differentially expressed for the effect of diet, 7.8% (978 genes) for the effect of the interaction between diet and tissue type (LRT, *P*_adj._ < 0.05). Interestingly, we observed similar patterns of gene expression relative to the C diet, in the DR and HS treated flies, a response likely reflecting diet component ratios. Hierarchical clustering identified 21 robust gene modules showing intra-modularly similar patterns of expression across diets, all of which were highly significant for diet or diet-tissue interaction effects (false discovery rate, FDR *P*_adj._ < 0.05). Gene set enrichment analysis for different diet-tissue combinations revealed a diverse set of pathways and gene ontology (GO) terms (two-sample t-test, FDR < 0.05). GO analysis on individual co-expressed modules likewise showed a large number of terms encompassing a large number of cellular and nuclear processes (Fisher exact test, *P*_adj._ < 0.01). Although a handful of genes in the IIS/TOR pathway including *Ilp5*, *Rheb*, and *Sirt2* showed significant elevation in expression, known key genes such as *InR*, *chico*, insulin peptide genes, and the nutrient-sensing pathways were not observed.

**Conclusions:** Our results suggest that a more diverse network of pathways and gene networks mediate the diet response in our population. These results have important implications for future studies focusing on diet responses in natural populations.

## Background

Individuals can withstand changing nutritional conditions by flexibly adjusting the allocation of resources to competing life history traits, allowing populations to adapt and thrive. Individual ability to partition available nutrients and optimize fitness gains requires complex cooperation at multiple levels of functional and structural organization in tandem with prevailing conditions dictating nutrient availability. Changes in diet are associated with many phenotypic changes across the tree of life. For example, in many metazoan species, moderate nutrient limitation extends lifespan and delays age-related physiological decline [1–4]. In fluctuating resource conditions, this effect, in which the individual often shifts nutrients away from reproduction and towards somatic maintenance and repair may be adaptive, ensuring survival in bad conditions and reproduction when good conditions return [5, 6]. On the other hand, constant dietary excess such as diets high in sugar, promote hyperglycemia in many genetic backgrounds, accelerate the rate of aging, and reduce lifespan [7–10].

A large and growing body of literature points to endocrine pathways being involved in nutrient perception and balance in order to coordinate organismal response to diet change. Nutrient sensing pathways are associated with aging and longevity from yeast to mammals [11–14], reviewed in [15–19]. The insulin/insulin-like signaling (IIS) together with the target of rapamycin (TOR) are among the most studied pathways. These pathways jointly regulate multiple metabolic processes affecting growth, reproduction, lifespan, and resistance to stress [20–22]. In insects, IIS/TOR signaling determines body size by coordinating nutrition with cell growth, and steroid and neuropeptide hormones to cede feeding when the target mass is attained [23]. Mutations, including experimental gene knockouts, that reduce IIS/TOR signaling reduce growth and reproduction, and increase stress resistance and lifespan [12, 24, 25], and apparently coordinates nutrient status with metabolic processes. For example, lack of nutrients blocks insulin production [26] and mimics the effects of a down-regulated IIS/TOR [27], while a hyperactivated IIS/TOR pathway can exclude the necessity for nutrients [27]. Fruit flies raised on excess sugar diets as larvae become hyperglycemic, fat and insulin resistant, and show increased expression of genes associated with gluconeogenesis, lipogenesis, β-oxidation, and FOXO effectors [8, 9]. Additionally, modulating TOR signaling slows aging by affecting downstream processes including mRNA translation, autophagy, endoplasmic reticulum stress signaling, and metabolism (reviewed in [28]).

Specific examples on the role of nutrient sensors abound in literature. Briefly, the forkhead transcription factor *foxo* in *Drosophila melanogaster (D. melanogaster)* and *foxo* orthologs in the nematode *Caenohabditis elegans* (*daf-16*) and vertebrates (*FoxO*) is the main transcription factor target of IIS/TOR, and is required for lifespan extension by a reduced IIS, reviewed in [18]. An activated *foxo* represses production of insulin-like peptides (ILPs) which in turn reduces IIS signaling [29, 30]. In a related mechanism, resveratrol-mediated activation of sirtuin genes mimic the effect of dietary restriction and promote lifespan in many metazoan species [1]. For example, in the cotton bollworm *Helicoverpa armigera*, *Sirt2* extends lifespan by its role in cellular energy production and amino acid metabolism [31, 32]. Further, the regulation of appetite which has a major effect on plastic nutrient allocation (reviewed in [33]), depends on leptin signaling together with the AMP-activated protein kinase (AMPK), influencing nutrient intake and subsequent production of ILPs [34–36]. Lastly, the hormones ecdysone and juvenile hormone also bear on the IIS to regulate ovary size and influence dispersal-reproduction trade-offs in *D. melanogaster* and sand crickets, *Gryllus firmus*, respectively *[21, 37–40]*, reviewed in [33]. In spite of these and other examples that demonstrate the effect of genetic variation on the metabolic response to nutrition, the underlying genetic basis diet effects in natural populations remain elusive [41].

Much of the current focus on how endocrine mechanisms affect phenotypic response to nutrition proceed in one-gene-at-a-time knockout strategies to elucidate function. This approach has been informative, largely in model species, but also supported to some extent in wild species. Endocrine pathways have been shown to affect plastic and adaptive resource allocation in wild *D. melanogaster [42, 43]*, sexual selection of horn size in rhinoceros beetles [44], sex-specific mandible development in staghorn beetles [45, 46] and morph determination in wing dimorphic sand crickets [38, 47–49], leading to the conclusion that endocrine pathways mediate the evolution of resource allocation strategies [50–52]. However, natural populations have not consistently revealed these same genetic mechanisms [53–56] suggesting that large effect studies in mutants capture only the tails of effect distributions that occur in the wild [57], or that different mechanisms overlapping with endocrine pathways may be involved [58, 59], reviewed in [33]. This disconnect means that our understanding of the specific genetic mechanisms that govern the response to diet in natural populations remains limited. In particular, there has been little characterization of the genome-wide transcriptional changes associated with diet. However, high-throughput genome-wide studies have become increasingly feasible, and generate high dimensional multi-level data from high replication experiments, allowing for a more comprehensive picture of these processes, and providing the opportunity to integrate data types such as QTL and gene expression studies.

In this study, our goal is to understand the transcriptional response in different nutritional environments in an outbred multiparent population. We use a synthetic population derived from the *Drosophila* Synthetic Population Resources (DSPR) [60, 61], and analyze RNA-seq data sequenced from pooled samples of female *D. melanogaster* exposed to multiple diet conditions differing in the proportion of protein and carbohydrate sources: dietary restriction (DR), control (C) and high sugar (HS). We profile gene expression for three tissues: heads (H), bodies (B) and ovaries (O), in high replication, and ask:

1. How does gene expression change in response to nutritional environment?
2. What specific biological processes and pathways are significantly perturbed by diet treatment?
3. Which sets of genes show similar expression patterns across diets and tissues, and what biological processes are involved in these specific patterns?

## Results

### Global expression patterns

We sequenced total RNA from 54 samples comprising 18 samples each for DR, C, and HS diet treatments (Fig. 1). In each diet treatment, samples for three tissues (head, body and ovary) were collected in six replicates. A summary of transcript levels across diets and tissues are shown in Figure 2. A total of 35,572 transcripts were obtained, out of which 18,678 transcripts remained for further analysis after removal of transcripts with a variance across samples of less than one [62]. Overall expression levels were generally consistent across diet treatments and tissues (Fig. 2a). One sample (bodies, B2) in the DR treatment showed slightly lower median expression compared to the rest, but was similar enough to the others and was retained in the analysis.

**Figure 1:**
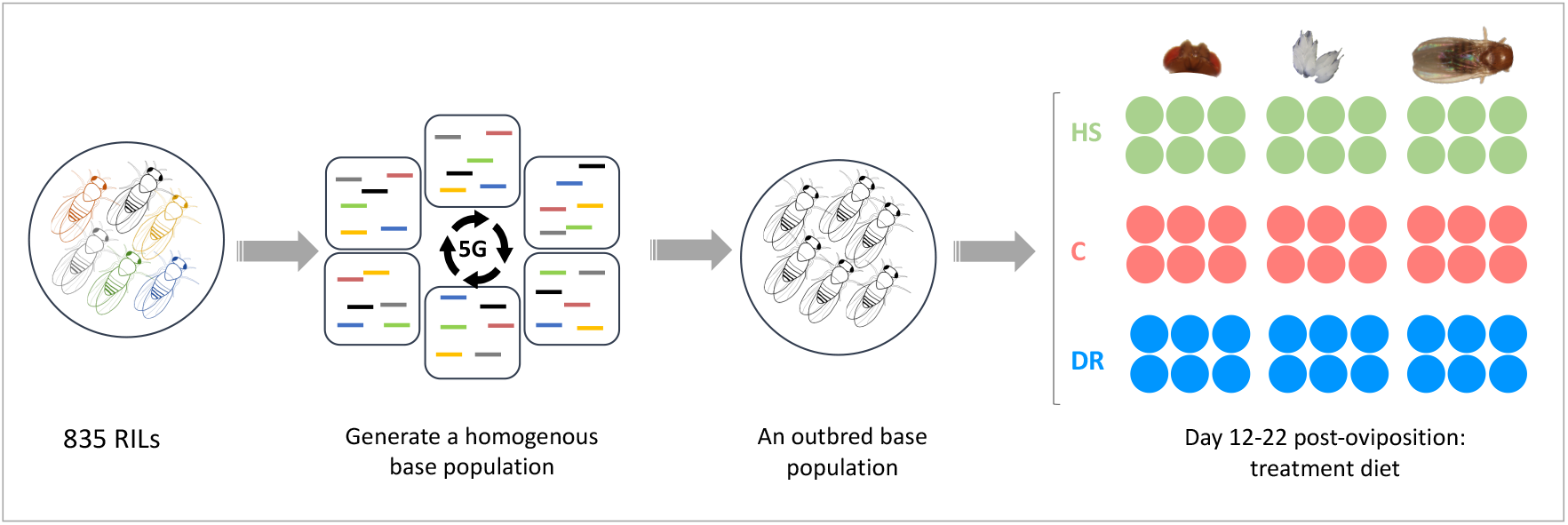
Study design. Flies drawn from 835 RILs of the DSPR were bred together for 5 generations to create an outbred panel. Eggs were collected from this homogenous population and resulting flies reared on dietary restriction (DR), control (C) and high sugar (HS) diets in six replicates for 10 days from day 12 post-oviposition. Heads, ovaries and bodies were dissected over 100 female flies from each treatment replicate for total mRNA extraction.

**Figure 2:**
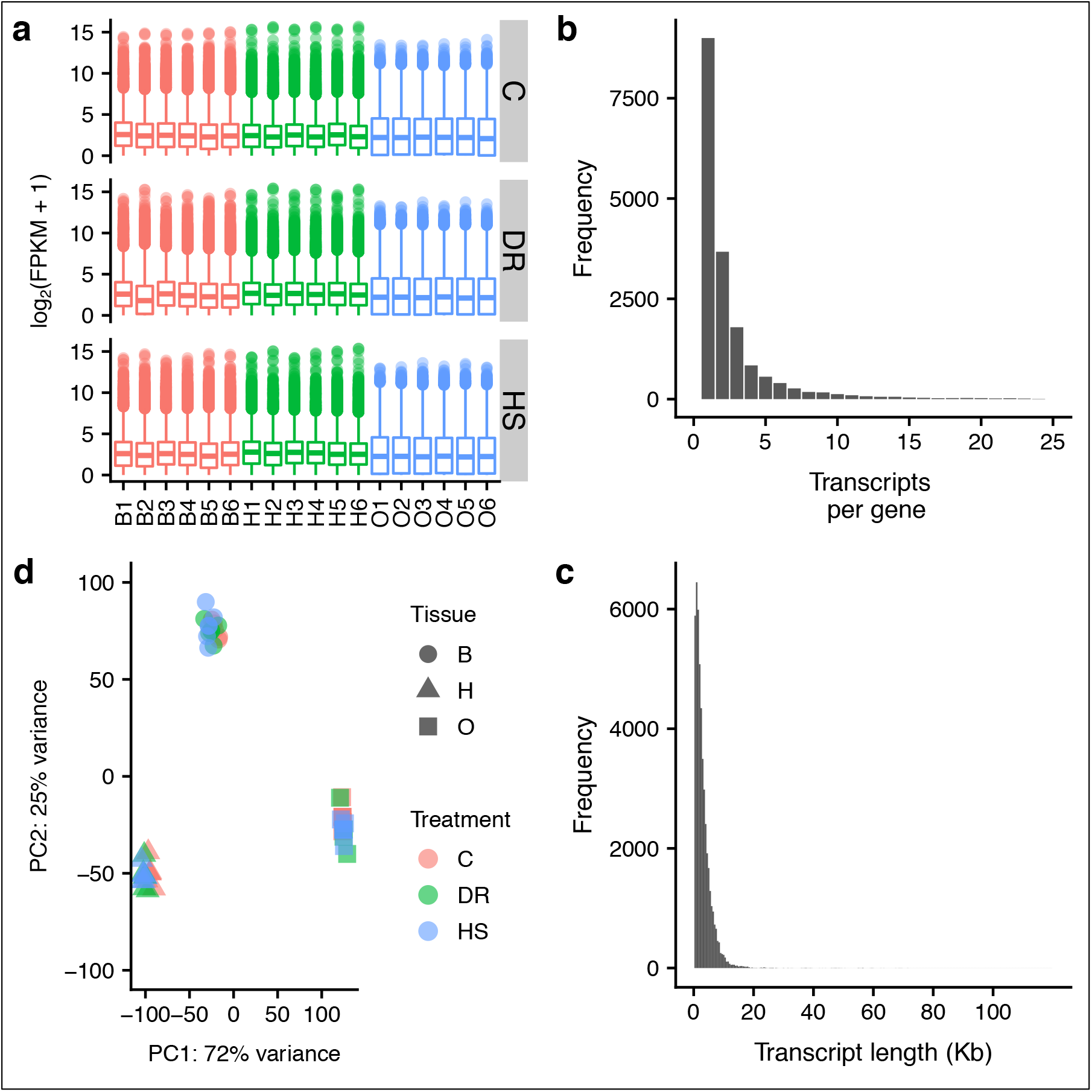
Global patterns of gene expression in 54 RNA samples. **a.** Transcript abundance as log2(FPKM + 1) for each sample (x-axis: bodies, B1 - B6; heads, H1 - H6; and ovaries, O1 - O6), and diet C, DR, and HS. **b.** Distribution of transcripts per gene - the number of transcripts associated with each gene symbol. **c.** Distribution of transcript sizes - based on transcript models of annotated genes. FPKM - Fragments Per Kilobase Million. **d.** Principal components analysis (PCA) to visualize the overall effect of diet and tissue. Two dimensions are shown (PC1 and PC2).

To assess global expression patterns over tissues and diets we performed principal components analysis (PCA) on all samples using an expression matrix from which batch effects had been removed (Fig. 2b). A similar figure prior to batch removal is shown in Additional file 1. As expected, tissue effects strongly dominated variance in the first two components which jointly accounted for 97% of the total variance. PC1 which explains 72% of the variance in expression presents non-overlapping separation of tissue expression, although body and head expression appear somewhat similar compared to the ovaries. PC2 (25%) distinguishes expression in bodies from that in heads and ovaries.

### Differential gene expression in response to diet

We used DESeq2 to quantify differential gene expression in head, ovary and body samples obtained from adult flies held on C, DR, and HS diet treatments. We obtained lists of genes significantly differentially expressed due to the main effect of diet. After filtering out genes with a low overall count, a total of 12,614 genes remained in the experiment based on which we report all subsequent results. Of these, 2,475 genes (19.6%, Additional file 2) were differentially expressed in response to diet treatment, and 978 (7.8%, Additional file 3) for the interaction between diet and tissue (LRT, *P*_adj_ < 0.05). The overall expression differences are visualized for each tissue and diet pair in Figure 3. Overall, relative to the C diet, many genes in all organs were expressed in the same direction in the DR and HS diets, meaning that the genes that have increased expression in the DR diet tend to also have increased expression in HS, and vice versa. This general trend is apparent in all tissues suggesting a similar change in global transcription pattern in response to both the DR and HS diets relative to the C diet, despite their very different compositions by weight and subsequently their caloric content. Further, the 2,475 DEGs for the main treatment effect were distributed across all diet-tissue combinations making it challenging to narrow down to a smaller list for further examination (Fig.4).

**Figure 3:**
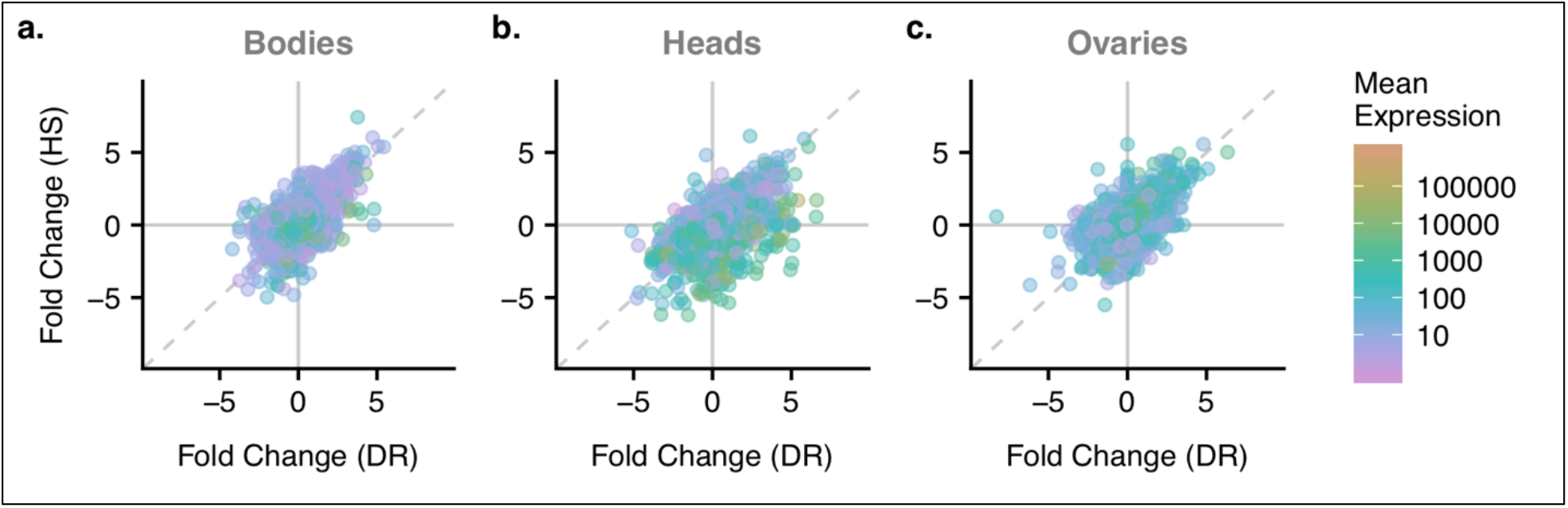
Comparison between DR and HS fold changes. Horizontal and vertical lines at 0 show when gene expression in the two diets is the same relative to the C diet. Diagonal dashed line is the 1:1 line. Points in the quadrants above 0 for one diet and below 0 for the other are genes that trend in different directions in the HS vs. DR diet relative to C (top-left and bottom-right). Points falling above the 1:1 line in the top-right quadrant and below the 1:1 line in the bottom-left quadrant show a similar effect in the HS diet as in the DR diet. Points are colored according to their mean expression.

**Figure 4:**
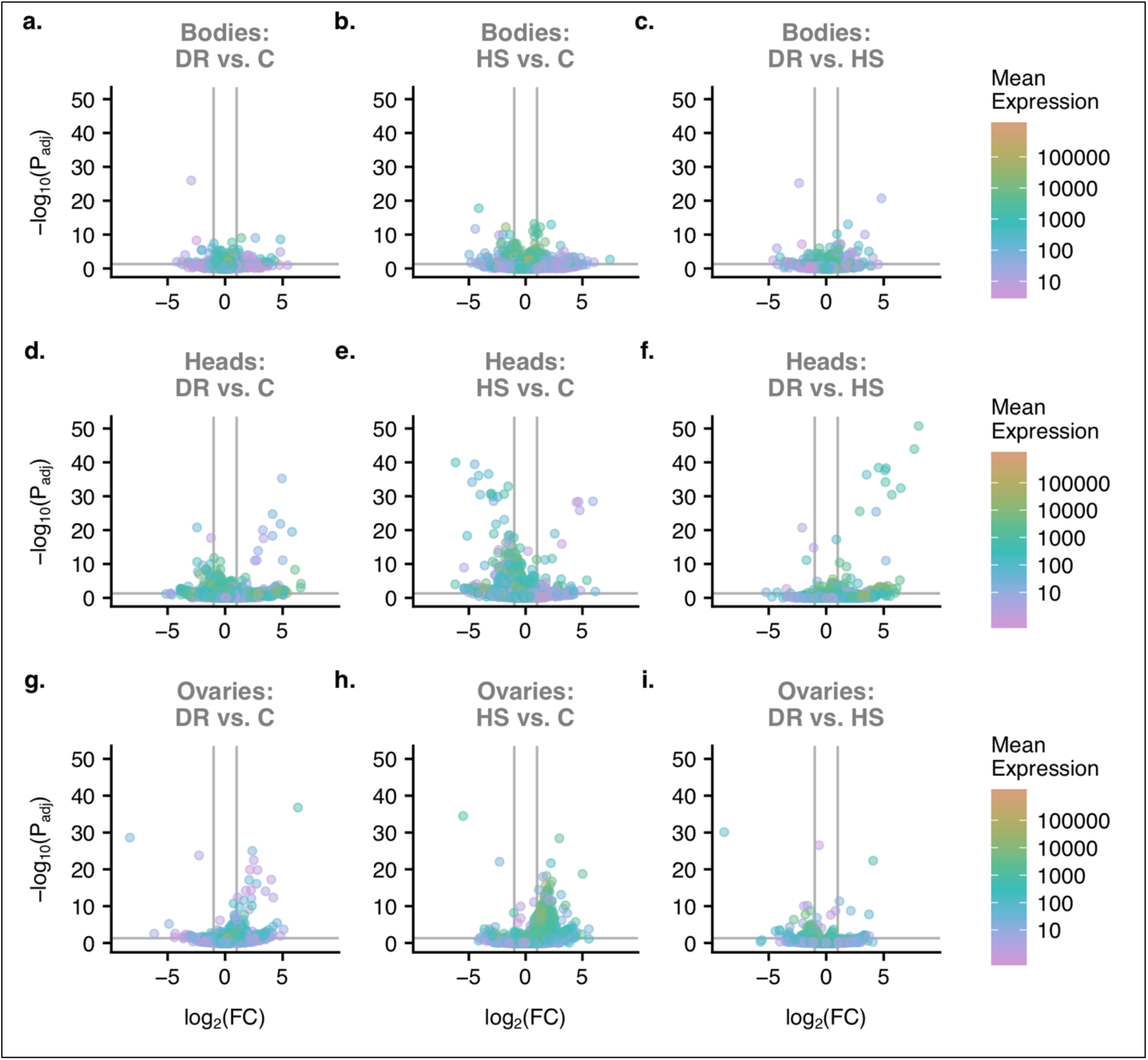
Volcano plots (**a - i**) for differentially expressed genes showing genes with large fold changes that are also statistically significant. Horizontal lines indicate −log_10_(P.adj) = 0.05, and points above the line represent genes with statistically significant differential expression. Vertical lines differential expression with the value of log_2_ fold change of 1 (i.e. absolute fold change = 2) and FDR = 0.05. Upregulated and downregulated genes are on the right side and left side of the vertical lines, respectively, and statistically significant genes are above horizontal lines. Rows in the panel top to bottom are bodies, heads, and ovaries; columns left to right are DR vs C, HS vs C, DR, vs HS; color of points represent log_10_ of base mean expression.

### Gene set enrichment analysis

We performed gene set enrichment analysis (GSEA) on the significantly differentially expressed genes (i.e. 2,475 DEGs) for the main effect of diet, using the fold changes for each diet-tissue combination to identify pathways and gene sets which were significantly perturbed relative to all DEGs in the model. Of these pairwise comparisons, only DR versus HS in bodies and DR versus C in bodies showed evidence for significantly enriched gene sets/pathways at an FDR *P*_adj._ < 0.05 (Benjamini & Hochberg procedure). We identified four pathways showing gene set level changes for bodies in DR relative to HS: Metabolic pathways (two-sample t-test, mean change = 5.38, FDR = 2.94e^−06^), Carbon metabolism (two-sample t-test, mean change = 3.31, FDR = 2.26e^−02^), Oxidative phosphorylation (two-sample t-test, mean change = 2.95, FDR = 4.52e^−02^), and Protein processing in endoplasmic reticulum (two-sample t-test, mean change = 2.83, FDR = 4.52e^−02^, Additional file 1). Notably, metabolic pathways (dme01100), which was most significantly enriched, is a large group of pathways in the KEGG database (https://www.genome.jp/kegg-bin/show_pathway?dme01100). At the default threshold (FDR *P*_adj._ < 0.1) in GAGE, ten more pathways appeared for DR relative to HS in bodies (Additional file 4). These additional pathways encompass three main metabolic themes: carbohydrate, amino acid and protein, and drug/xenobiotics. For the comparison of DR vs C in bodies, oxidative phosphorylation (dme00190) was significantly enriched (two-sample t-test, mean change = 3.2, FDR *P*_adj._ = 7.36e^−02^).

Further, we examined GO term gene set enrichment for biological process (BP) to capture significant diet-related differences occurring below the level of pathway. Four terms were enriched at an FDR *P*_adj_ < 0.01. Small molecule metabolic process was enriched for the DR vs HS comparison in bodies (mean change = 4.49; P_adj_ = 5.84e^−3^). Cell communication (mean change = 5.10; P_adj_ = 1.83e^−4^), signaling (mean change = 5.06; P_adj_ = 1.83e^−4^), and signal transduction (mean change = 4.56; P_adj_ = 1.37e^−3^) were all enriched for the HS vs C comparison in heads. At an FDR *P*_adj._ < 0.05, 41 unique enriched terms were observed, of these, 34 terms were enriched for HS relative to C diet in heads (Additional file 4). These terms highlighted a broad range of themes including signaling, metabolism, growth, cytoskeleton, gene expression and development. Three terms were enriched for HS relative to C in bodies, including cell communication, signaling, and system process. The remaining six terms were all for the HS diet relative to DR in bodies, all within one theme of metabolism (acid, small molecule, carbohydrate). No terms were enriched for the comparisons in ovaries. To understand broader inclusive processes represented by these GO terms, we evaluated our list for ancestral terms using QuickGO (EMBL-EBI https://www.ebi.ac.uk/QuickGO/). Nine ancestral terms at the same hierarchical level immediately below category BP were observed (metabolic process, biological regulation, cellular process, localization, response to stimulus, cellular component organization, multicellular organismal process, growth, and developmental process). Among these, metabolic process, cellular process, and developmental process had the most connections to child terms. Our GSEA analysis therefore highlights multiple pathways and biological processes triggered by diet changes, especially in bodies and heads, and encompassing broad themes from metabolism to signaling to homeostasis, but none of the canonical nutrient sensing pathways such as IIS/TOR and FOXO signaling pathways. Notably, our results do not show particular enrichment of diet-associated terms in ovaries, at least for biological processes.

### Diet-induced gene co-expression

Next, we asked how diet treatment affected gene co-expression across tissues and diets, to identify sets of genes showing similar changes across diets and tissues. To identify co-expressed genes (or modules) we used gene expression profiles with batch effects removed. Setting the minimum module size to 30 genes, a total of 31 modules were detected (range gene number 39 - 3,240), with 1,049 unassigned genes (grey module). After merging highly similar modules (i.e. eigengene correlation *r* > 0.9, see methods), 21 modules were identified with an additional module holding all unassigned genes (Fig. ADDITIONAL FILE 4).

To appreciate module-level effects of diet and tissue on coexpression, we visualized eigengene expression across diets (Fig. 5, Additional file 5). It is clear from these plots that some modules showed greater diet by tissue interaction effects than others (e.g. *e*, *f*, *m*, *q*, *s* and *v*). These modules show either reduced or increased expression for one or two tissues in one or two diets. To gain better insight into these intra-modular effects of diet and diet-tissue interaction, we fit an analysis of variance model (ANOVA) to module eigengenes. For the main effect of diet, all modules turned up significant (FDR *P*_adj._ < 0.05), except modules *c* (Fig. 5). Similarly, for the effect of the interaction between diet and tissue, all modules showed a significant effect (FDR *P*_adj._ < 0.05), except module *a*.

**Figure 5:**
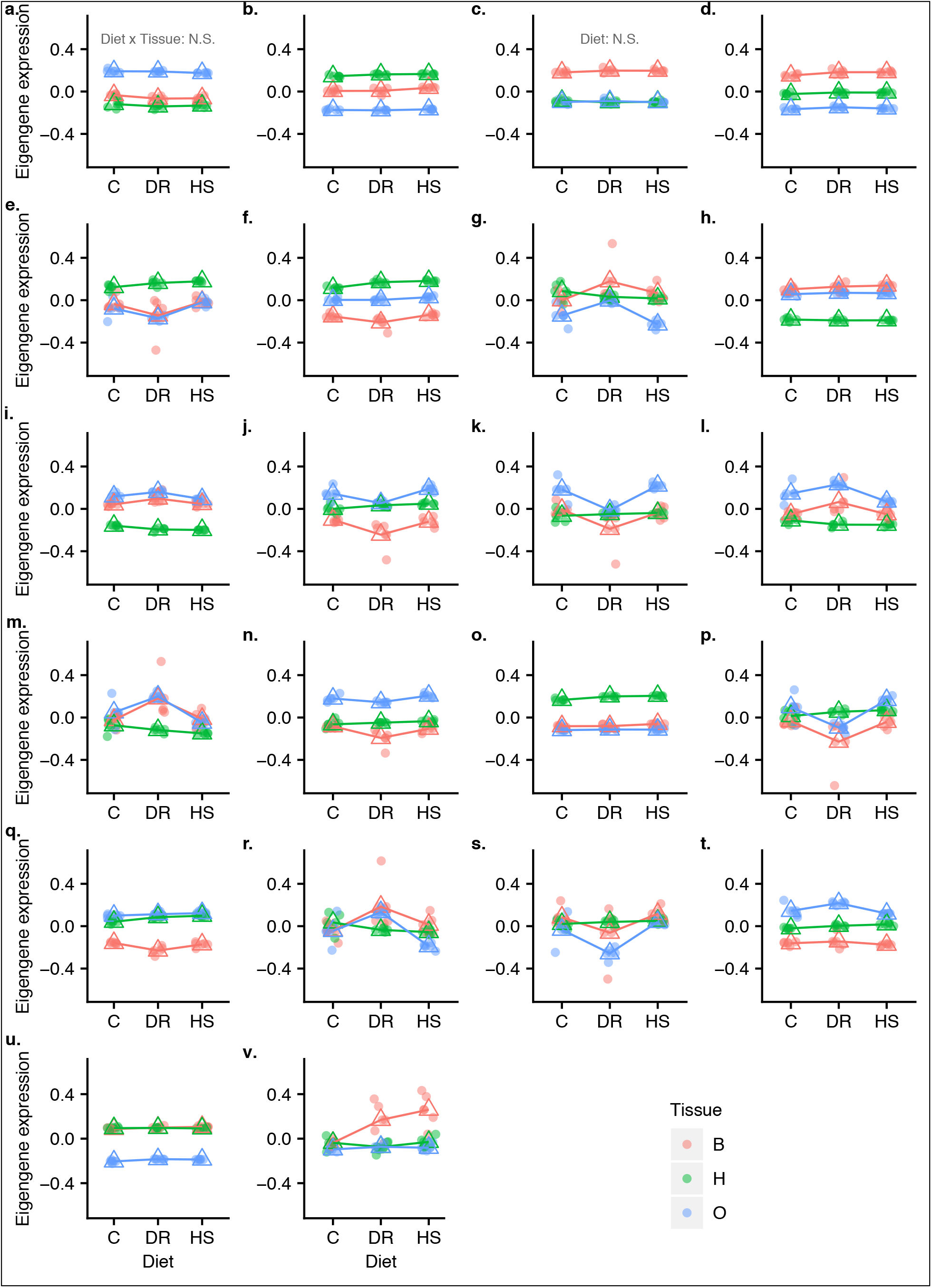
Eigenegene expression across diet treatment for each module *a - v* identified in hierarchical clustering. Color scheme represents the three tissues; each filled circle represents a sample; the open triangle marks the mean eigene expression for a given diet in a given tissue. In all cases except those noted on (a) and (c), the main effect of treatment and the tissue by treatment interaction are significant.

Focusing on the modules showing a statistically significant interaction effect, and divergent expression profiles in one or more diets for a given tissue (Fig. 5), several distinct patterns became apparent: 1) generally reduced expression in the DR diet for ovaries and bodies unlike the rest of diets (Fig. 5*e*,*f*, *k* and *s*), 2) increased expression in the DR diet for ovaries and bodies (*i*, *m*), 3) elevated expression in bodies in both DR and HS diets (*v*), and 4) different responses in all three diets (*g*, *r*). An attempt to isolate specific diet-tissue combinations driving the interaction effect using *post hoc* tests revealed large numbers of highly significant combinations. We therefore explored the modules further via functional enrichment to identify the processes driving these coexpression patterns.

We conducted functional analysis on all modules to identify enriched GO terms (Bonferroni corrected enrichment *P* values, Additional file 6). Of 12,614 entrez identifiers in our experiment, 10,334 mapped in current GO categories (see methods), and therefore used as a background list for enrichment analysis in WGCNA. A large number of terms were obtained across CC, MF and BP categories: 658 terms (*P* < 0.01), and 791 terms (Bonferroni corrected *P* < 0.05) (Additional file 6).

A visual inspection of enriched terms in the 21 robustly assigned modules confirmed a large diversity of highly significantly enriched biological processes in most modules, ranging from nuclear processes, to membrane and cytosolic processes; from structural to signaling and immune response processes; and from pigmentation to homeostatic processes (Additional file 6).

The first module (Fig. 5*a*) which included 2,956 showed 291 GO terms (Bonferroni corrected, *P*_adj._ < 0.01), and had the most significantly enriched terms (i.e. > 60 terms ranged between *P*_adj._ < e^−156^ − <e^−15^). This module was characterized by greater eigengene expression in ovaries compared to heads and bodies, although the diet effect was subtle but significant. Nuclear and intracellular organelle processes including gene expression, and RNA processing were key tissue (ANOVA, *P* < 2e-16) and diet (ANOVA, *P* < 0.002) effects independently regulated (i.e. no interaction effect). With reference to the trends described above (Fig. 5), those modules showing generally reduced expression in the DR diet for ovaries and bodies (*e*,*f*, *k* and *s*), are associated with biological processes including signaling (*e*, *P*_adj._ <1.1e^−10^), cellular component organization (*k*, *P*_adj._ <5.8e^−09^), nervous system development (*f*, *P*_adj._ <1.3e^−14^), signaling and protein localization on Golgi apparatus (*s*, *P*_adj._ <3.0e^−06^). Interestingly, expression increase in DR in bodies and heads compared to ovaries is related to ubiquitin-dependent proteolytic processes in the proteasome (*i*, *P*_adj._ <1.8e^−08^), and cytosolic vesicle transport/mitochondrial activities (*m*, *P*_adj._ <8.9e^−156^). Module (*v*, *P*_adj._ <1.1e^−21^) was interesting because bodies show monotonic increase in expression from C to DR to HS, a trend that may relate to the GO term chitin-based cuticle structure development (*P*_adj._ = 5.78e^−30^), indicating cuticular remodeling in stressful diets (DR and HS), presumably to accommodate gain or loss of body mass.

Analysis of our modules therefore revealed a large number of biological processes (BP), molecular function (MF) and cellular components (CC) (Additional file 6), suggesting that response to diet changes in natural *D. melanogaster* involves a multi-system response rather than one or a few signaling pathways that can be very different in different tissues.

### Previously implicated pathways

Several pathways have been well-studied in the context of diet-induced phenotypic changes (see Background). We specifically examined these pathways in order to characterize the transcriptional response. We obtained a precomputed list of genes for key metabolic pathways (fbgn_annotation_ID_fb_2019_02.tsv): IIS (55 genes), TOR (152 genes), and FOXO (110 genes, which includes the *Sir* genes). Of 317 pathway genes, 47 were significantly differentially expressed for the diet effect (Additional file 8). However, our GSEA results presented above did not show pathway level enrichment of any of these pathways as defined in KEGG Pathway Database (https://www.genome.jp/kegg/pathway.html), although as mentioned above, a handful of member genes were differentially expressed. These genes that were differentially expressed however, showed mixed modular membership in our clustering results. Similarly, GO analysis on module genes did not enrich categories specifically including ‘insulin’, ‘TOR’ and ‘FoxO’ in the term. From these results we could not link any of these pathways with particular modules within our data. Importantly, we did not find evidence for significant whole pathway perturbation of IIS/TOR and the downstream FOXO signaling in our co-expressed modules. Expression of a sample of genes frequently associated with nutritional effects is shown in Figure 6. With the exception of *Ilp5* and *Sirt2*, these were not all not significantly differentially expressed in this study.

**Figure 6:**
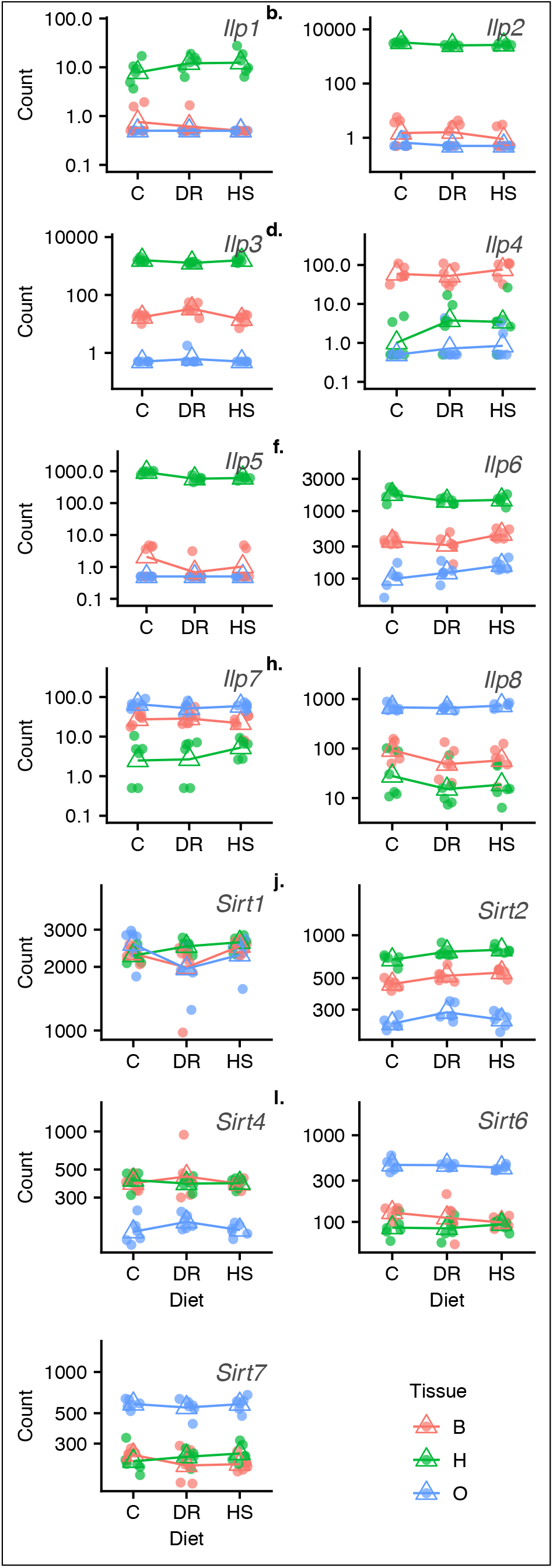
Expression of a sample of genes that are frequently associated with nutritional effects in may model organisms. Only *Sirt2* and *Ilp5* were significantly differentially expressed in this study.

### Parsing previously identified QTL for the response to diet

A useful application of genome-wide expression data is to identify possible regulatory variants underlying QTL. In a previous study, we used a multivariate approach to identify a global expression QTL for the response to diet treatment of 52 genes in the IIS/TOR pathways [63] that we hypothesized was influencing the expression of many of the genes in these pathways. After performing a discriminant function analysis predicting diet (DR or C) from expression measured on whole flies of 52 genes in the IIS/TOR pathways, we mapped the difference in discriminant function to identify this eQTL. The eQTL interval, as defined by the Bayesian credible interval, contains 327 genes, making it challenging to narrow to possible candidates. We therefore searched for differentially expressed genes identified in this study that fall within this eQTL region of interest. Of these, we find 49 genes are differentially expressed in the different diet treatments and 13 show a diet by tissue interaction, with 5 of these genes showing both a main treatment effect and an interaction effect (i.e. *Odc1*, *Dgat2*, CG12822, CG12159, and *Obp44a*, Additional file 7). The patterns of expression across tissues for this set of genes is shown in Figure 7. While our expression results don’t allow us to narrow to a single candidate, they do significantly reduce the list and provide detailed expression data for the possible causal genes.

**Figure 7:**
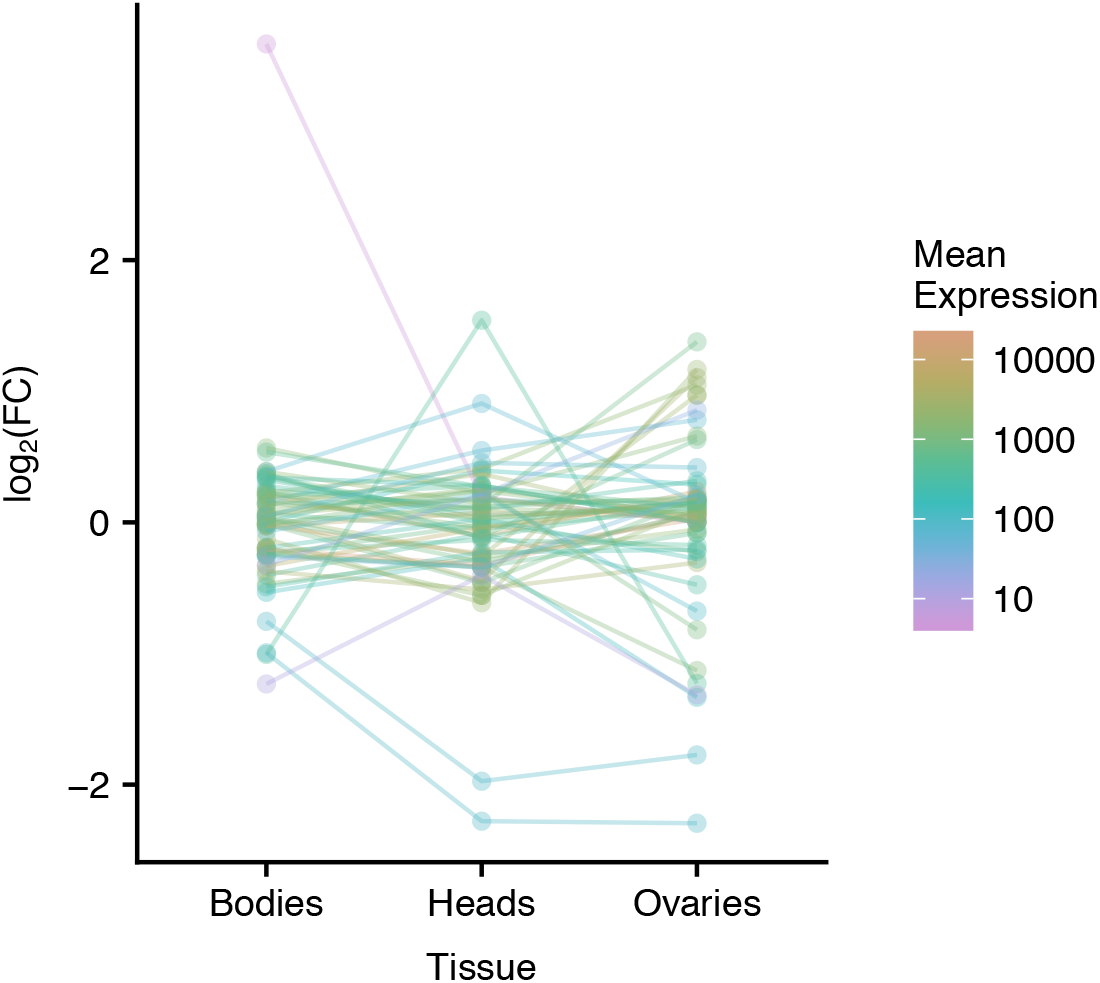
Individual gene plots representing a sample of key genes hypothesized to underlie canonical nutrient sensing pathways, particularly the IIS/TOR and sirtuin deacetylases which are thought to regulate growth, reproduction, lifespan and stress resistance in many species including *D. melanogaster*. *Ilp* - insulin peptide gene, *Sirt* - sirtuin gene. There are 7 insulin peptides and 5 sirtuins known in *D. melanogaster*.

## Discussion

We sequenced pooled RNA samples from a three-diet by three-tissue by six-replicate experiment of outbred mated female *D. melanogaster* in the DSPR. Our aim was to understand diet-induced patterns of gene expression influencing plastic nutrient allocation in different diet conditions in a multiparent population resource. Our results suggest that: 1) global expression patterns are dominated by tissue and diet-tissue interaction effects, while the effects of diet alone are subtle but significant, 2) patterns of gene expression are generally similar in low-protein and carbohydrate-rich diets relative to the control diet, 3) multiple pathways, co-expressed gene modules, and biological processes are invoked that affect transcription in different diet conditions, especially in the head and body tissues, and 4) expression results did not suggest a single regulatory variant underlying QTL, but narrowed down to a few possible causal genes. Overall, our results suggest that multiple networks are involved in phenotypic changes in response to nutrient availability, rather than just a few key genes. We advocate a broader, genome-wide approach to studying the genetic mechanisms underlying diet effects on phenotypic change.

It is established that nutrient signaling pathways and hub genes in those pathways play a crucial role in how organisms adjust to changing conditions in availability and quality of nutrients to optimize fitness traits [50–52]. Analysis of differential gene expression in a population presented with diets differing in nutritional richness provides an ‘omic’ alternative to study intermediate processes that connect genetic architectures to phenotypic outcomes such as allocation patterns. In fruit flies, studies typically focus on whole-body or head tissue transcription (e.g. [64]); one or a few gene pathways known to affect diet responses at a time (e.g. [63]); one or two diet manipulations (e.g. [64–66]), but scarcely integrate over more than two organs and conditions at a time, or explore expression outside a few gene pathways. Further, despite costs trending downwards recently, sequencing of more highly replicated experiments remains unaffordable for many laboratories. By and large, studies in model organisms focus on genes in a few endocrine pathways, so called nutrient sensors, as critical players in coordinating growth, reproduction, stress resistance and somatic maintenance responses to changing diet conditions. Components of the IIS/TOR, growth, and ecdysone hormones; and sirtuin deacetylases are deemed some of the major players in this respect. Our results suggest an expanded scope of mechanisms underlying flexible responses to nutrient limitation (DR studies) or oversupply (high sugar and high fat diet studies) in natural populations, which has also been suggested by [67–69] and reviewed by [58, 70]. We discuss our major results and their implications below.

First, we observed a large global effect of tissue type and a more subtle, but significant, effect of diet treatment. Previous studies in flies have also found relatively small effects of diet on transcription. Previously, we characterized the genetic basis of standing genetic variation for 55 genes of the IIS/TOR pathway following treatment with the same diets used here [63], and found only small changes in gene expression associated with diet treatment although most of those genes were differentially expressed. Similarly, Reed et al, [71] measured transcriptional and metabolomic changes for 20 inbred lines (North Carolina and Maine population) of *D. melanogaster* treated with four diets varying in sugar and fat content and observed a small dietary component in gene expression profiles, with much larger contributions of genotype by diet interactions. Musselman et al. [72] investigated expression differences in *D. melanogaster* fed with two different forms of sugar and found small but significant changes in gene expression. Overall, diet seems to produce fairly small magnitude changes in expression in many genes across the genome, which in concert presumably can lead to large phenotypic changes.

Secondly, comparisons between DR and HS diets relative to C revealed a similar pattern of expression. This result mirrors our earlier finding in an eQTL mapping experiment using DSPR lines in which gene expression in DR and HS relative to C generally trended in the same direction [63]. This result is in spite of the fact that the DR and HS diets lead to very different outcomes in median lifespan and fecundity in our population [63, 73]. Nutritional geometry studies which measure traits in a series of concentrations of liquid media suggest that traits such as lifespan and reproduction (which differ significantly across our diets) are influenced primarily by the diet protein to carbon (P:C) ratio, not its caloric content (e.g., [8, 74, 75]. Thus, calorie limitation alone does not drive phenotypic patterns in these studies. While our HS diet has a high concentration of sugar per liter of food, the P:C ratio is a lot lower (i.e. ~1:2 yeast to sugar ratio). Lifespan and fecundity in nutritional geometry studies are maximized at a much higher P:C ratio (i.e. 2- to 4-fold higher than our HS diet) [74, 75]. Thus, our results are consistent with similarity in expression pattern between DR and HS. It is also possible that diet macromolecules serve only as a cue that lead to optimal allocation of resources in natural populations. Thus, it is possible for expression measures to trend in the same direction in DR and HS treatments. Further, Dobson et al. [76] found that excess sugar diets in young adult flies inhibited *foxo* and reduced survival in middle and old age. While we did not measure lifespan and fecundity here, our data showed mixed pattern of *foxo* activation across diet tissue combinations, and these mapped to many coexpression modules which may be due to the relatively young age (i.e. 21 days) of our experimental flies.

Thirdly, an overall take away from our GSEA on the full dataset, and also on gene modules coming from hierarchical clustering was that diet effects could not be attributed to a particular genetic mechanism. GSEA highlighted metabolism, oxidative phosphorylation and protein processing at pathway level, but showed a broad spectrum of processes in GO term enrichment encompassing metabolism, cell signaling, structural development and organization, and defense. Similarly, GO analysis of gene modules yielded a broad range of biological processes. From both these analyses, IIS, TOR, and FOXO signaling were not significantly enriched. However, several genes had significantly reduced expression in the IIS/TOR (e.g. *Ilp5*, *Rheb*, *Atg1*, *Myc*, and *eIF4E1*) and significantly higher expression in FoxO downstream effectors (e.g. *AMPK, orct2*, *Gadd45, cdk2* and *p38*) in most DR-tissue combinations consistent with indicating canonical activity. With the exception of *Ilp5 [41]* however, undetectable differential expression of hub genes (such as *Ilp2, S6K*, *chico, InR, Akt1, Torc, Thor, and foxo*) suggest that diet induced effects may involve many more pathways/genes than have been traditionally studied in this context.

Evidence in *C. elegans* suggest that the worm ortholog of *foxo*, *daf-16* is not required in the DR response [77–80]. On the other hand, *sir-2.1* is a worm ortholog of the fruit fly *Sirt 2* (which was significantly differentially expressed in this study), and is required for lifespan extension in adult worms by diet deprivation was independent of the *daf-2*/insulin-like signaling [80]. In *D. melanogaster*, similar evidence is emerging that suggest that *foxo* is not required for the response to DR [81], but is involved in the normal response to DR [82]. When *dFOXO* was removed, DR treatment still resulted in significant lifespan extension in null flies [81]. Another study testing a novel DR assay in *C. elegans* found that DAF-16, but not DAF-2 (the worm *lnR*) was required when DR was performed on solid media, and concluded that AMPK-FOXO signaling resulted in lifespan extension on solid food [83]. Our data provides further evidence supporting these studies in suggesting a broader mechanism in which IIS, TOR and FOXO play a role, but in concert with other pathways.

A potential limitation of our study is the heterogeneity in tissue types present in our samples, which may affect the level and nature of gene expression [84]. For instance, assuming fewer cell types are available in the ovaries or head samples compared to body samples, we could expect the range of biological processes triggered by nutrient levels to scale with cell types to some degree. Further, DR and HS protocols vary tremendously across laboratories which can result in different studies detecting only subsets of the gene network distribution which responds to nutrient change [81]. Patterns of phenotypic expression (fecundity and lifespan) and patterns of gene expression have held stable across several studies using the same set of diets in our population, suggesting that the effects we find are biologically relevant in the DSPR. However, in addition to a broader view of the potential mechanisms causing phenotypic changes in response to diet, a broader consideration of different diets would also benefit the field.

## Conclusion

We have characterized the genome-wide transcriptional response to diet composition in multiple tissues in *D. melanogaster,* providing a comprehensive picture of potential genetic mechanisms underlying phenotypic changes. We found that the general pattern of expression was similar in DR and HS diets relative to C diet, probably reflecting specific nutrient ratios. In addition, we identified a large set of co-expression networks, pathways and gene ontology terms that were enriched in response to diet. Our data did not show enrichment of canonical nutrient sensing pathways and key genes, although some genes in those pathways were significantly perturbed. Our results therefore support the hypothesis that in natural populations, multiple gene networks and pathways are invoked to respond to environmental differences in nutrient availability, and not just a few pathways as it is often assumed. Our study has potential implications for future studies focusing on the effects of stressful diets in natural populations including our own. We therefore urge future studies to look beyond traditional genetic mechanisms governing diet effects.

## Methods

### Experimental population

We used 835 recombinant inbred lines (RILs) of the *p*B sub-population of the DSPR as source of our experimental population. Five young females that had mated intra-line were randomly selected from each RIL and placed in six custom-made cages each measuring 20.3 cm × 20.3 cm × 20.3 cm. Eggs were collected (twice on successive days) after 22 hours of oviposition in milk bottles (250 ml) containing 40 ml media. Resulting flies were introduced to six cages in large population sizes (i.e. > 2000 per cage) and allowed to mate for five generations (21-day cycle). To ensure a genetically homogenous base population was generated, eggs from each cage equally seeded each of the 5 other cages at each generation. We refer to these six cages as the base population. All life stages during generation of the base population were reared on a cornmeal-dextrose-yeast maintenance diet. Proportions of ingredients in our maintenance diet are presented in Additional file 9. Additional culture practices including equipment and supplies are described in [85].

### Study design

We used one random cage from the base population for phenotyping to generate RNA-seq data we present here. This population is referred to as the “synthetic population”. We adopted a factorial design comprising three dietary treatments, three tissues, and six replicates per each treatment-tissue combination. To obtain flies for each diet treatment, we placed two plates (100mm × 15 mm, Fisherbrand^®^) of maintenance diet in the synthetic population and allowed for 24 hours of oviposition, repeating egg collection three times. From the two egg plates, a thin slice of media bearing 50 - 90 eggs (visually estimated) was cut out and grafted to each of 60 new vials (25 mm × 95 mm, Polystyrene Reload, cat. no. 32—109RL, Genesee Scientific, USA) of maintenance food containing 10 ml media. At 12 days post-oviposition, eclosed flies were released into 9 cages, 3 cages for each of three diet treatments, such that treatments are equally represented across egg collection dates. For each cage, we used 20 vials to seed the cage with adult flies. Flies were held on treatment diet for 10 days, and provided with new food every 2 - 3 days. Each replicate was started on a new week to provide for time that would be needed to dissect a large number of flies per replicate later on. All flies were reared in a growth chamber at 23 °C, ≥50% relative humidity, and a 24:0 light: dark cycle, which are the typical maintenance conditions for the DSPR flies. Details of our study design are schematically shown in Figure 1.

### Experimental diets

We extracted RNA for sequencing from 22 day-old (post-oviposition) female flies held on three experimental diets adapted from Bass *et al*, [86] and Skorupa *et al*, [8]: C, DR, and HS for 10 days, as described above. These are all cornmeal-sucrose-yeast diets which differ in two ways: relative to C, DR contains 50% less yeast, and HS contains about seven times sucrose (Additional file 8). We have used these diets in past studies to map expression QTL in the DSPR RILs [63], and to estimate quantitative genetic parameters in an outbred DSPR population [73]. To prevent desiccation and preserve quality, diets were covered with multipurpose sealing wrap (Press’n Seal^®^) sealed and stored at 4 °C and used within 2 weeks of preparation. To prevent food degradation, individuals were moved to vials with fresh food three times per week.

### Sample preparation and sequencing

At day 22, we collected 100 - 104 females per diet treatment under mild CO_2_ anesthesia. We replicated this process six times over six days, thus creating six same-age sample batches with each batch including all three treatments (Table 2b). This yielded assays of at least 100 females per treatment diet (HS, C and DR), three tissues within each diet (heads, ovaries and bodies), biologically replicated 6 times. Our experiment therefore comprised 54 samples (18 per diet). We immediately dissected tissues from these females and promptly flash-froze them in liquid nitrogen and held them in a closed styrofoam box on dry ice before storage at −80 °C. Precision scissors (RS-5604, Roboz Surgical Instrument Co., Inc.) were used for fly dissection in a bath of pH 7.4, 1% PBS (Life Technologies™) containing 2 drops (80-100 μl) Triton x100 (SIGMA) under dissecting stereoscopes (Leica S6E and Leica M216) in 100 mm × 15 mm polystyrene petri dishes (Fisherbrand^®^). Scissors and forceps were rinsed with 70% ethanol and wiped dry with Kimwipes between samples. To minimize time-of-day effects, dissection was done across treatments (e.g. [C, HS, DR], [DR, HS, C], etc.) rather than one treatment at a time (e.g. [C, C, C], [HS, HS, HS], etc.).

We purified whole RNA from each of 54 samples using a protocol modified from Life Technologies TRIzol RNA extraction protocol. Tissues were homogenized in a tissue lyser using steel beads in 1 ml TRIzol reagent, and subsequent RNA extraction by following the TRIzol protocol. RNA quality was evaluated on a Nanodrop 2000 spectrophotometer (Thermo Scientific) and concentration on a Qubit 2.0 fluorometer (Invitrogen, Life Technologies). An additional clean-up step with a QIAGEN RNeasy Plus Mini Kit (Hilden, Germany) was used for gDNA elimination following manufacturer’s protocol to achieve high quality RNA for library preparation. After the RNA cleaning step, yields typically ranged between 400 - 800 ng/μl; and absorbance values (260/280 and 260/230) above 2.0 (Additional file 10). Fifty-four libraries were each prepared from 1.5 μg total mRNA, RNA integrity (RIN) ≥ 8.0 using Illumina’s TruSeq Stranded mRNA (poly A enrichment) and single-end read sequencing on Illumina NextSeq 500. Barcoded libraries were combined in a single 54-plex library, which we sequenced on three lanes of a NextSeq 1×75 bp run for an average of 23 million reads per sample. The resulting reads were trimmed of adaptor sequences and FASTQC was used to assess quality. Sequencing was performed at the University of Missouri DNA Core Facility.

### Data analysis

#### Read alignment and quantification of expression

We employed the ‘new Tuxedo’ suite analysis pipeline [62] for read assembly and transcript quantification. We mapped single-end reads to the reference genome of *D. melanogaster*, bdgp6_tran (ftp://ftp.ccb.jhu.edu/pub/infphilo/hisat2/data/, updated March, 2016,), using HISAT2 (version 2.1.0, Kim *et al*, [87]. Alignments were converted to BAM file format and runs combined using SAMtools [88]. Then, StringTie [62, 89] was used to assemble RNA-seq alignment into full-length transcripts, and to quantify levels transcript expression.

We intended our downstream analysis to focus on gene-level differential expression of known genes in the *D. melanogaster* transcriptome. Therefore we used the alternative workflow given by Pertea et al [62] skipping steps 3 - 5 in that protocol, (i.e. we skipped the individual assembly of each sample and the merge step, documented here http://ccb.jhu.edu/software/stringtie/index.shtml?t=manual#de, last accessed May 20, 2019). To do this, we ran `‘stringtie -eB’` directly on the output of `‘samtools -sort’`. In order to extract more gene (FBgn) ids from reference gene annotations into StringTie output, we ran a Perl post-processing script, ‘*mstrg_prep.pl*’ (Pertea, https://gist.github.com/gpertea/b83f1b32435e166afa92a2d388527f4b) that appends reference gene ids to the MSTRG gene ids used in StringTie. In order to perform differential expression analysis on row counts using DESeq2 [90], we processed the output from StringTie with a Python script ‘*prepDE.py*’ (Pertea M, http://ccb.jhu.edu/software/stringtie/index.shtml?t=manual#de) with the `-e` parameter to extract read count matrices (one for transcripts, one for genes) directly from the files generated by StringTie.

#### Controlling for batch effects

As described earlier, our samples were processed in six groups on separate days (all treatments and tissues represented equally in each) including setup, dissection, and RNA extraction and preparation (Additional file 11). We sought to control for this obvious batch effect and any unknown underlying batch effects. We therefore used surrogate variable analysis (SVA, [91] implemented in the R library svaseq [92]. We used the DEseq2 package [90] to obtain counts that are normalized for library size (i.e. counts divided by size factors). Eliminating genes with zero counts, we used the svaseq function comparing the full model with a known batch to a null model with batch only. A single surrogate variable (SV) was identified and included in all subsequent models.

#### Differential gene expression

Next, we estimated differential gene expression with DESeq2 for DR and HS treatment conditions relative to the C treatment. We fit two generalized linear models (GLM), with parameter `fitType = ‘local’` and only included genes with at least 10 reads in at least 2 samples. We compared the following GLMs:

1. Expression ~ SV + batch + tissue
2. Expression ~ SV + batch + treatment
3. Expression ~ SV + batch + tissue + treatment
4. Expression ~ SV + batch + tissue + treatment + tissue*treatment

where SV is the surrogate variable identified earlier. For all tests, we used a likelihood ratio test, comparing a more complex model [90] with a reduced model in the following way: 1) main effect of tissue: model 3 vs. model 2, 2) main effect of treatment: model 3 vs. model 1, 3) treatment by tissue interaction: model 4 vs. model 3. From these model comparisons, we identified sets of significantly differentially expressed genes using an FDR threshold of 0.05. To visualize and rank the genes we used the function `lfcShrink`, which performs shrinkage on log_2_(Fold Changes), which have been shown to produce better estimates. All log_2_(Fold Changes) reported here are the shrinkage estimated values using the `“normal”` estimator [90]. To identify global patterns of expression across diets and tissues, we performed a principal component analysis (PCA) on expression values before and after removal of batch effects.

#### Gene set enrichment in relation to diet

We considered our list of DEGs for the effect of diet treatment and used the R Bioconductor GAGE package [93] to infer gene sets and pathways that were significantly perturbed relative to all DEGs. Briefly, GAGE takes an expression matrix with log-based fold changes as input. It assumes that a particular gene set or canonical pathway (from literature or database) comes from a different distribution than the experiment-derived background list. Therefore a two-sample t-test is applied to account for the gene set specific variance and the background variance and derives 1) a global p-value using a meta-test on p-values from gene set comparisons, and 2) a FDR q-value adjustment using Using GAGE to access the Fly annotation database ‘org.Dm.eg.db’, we generated up-to-date KEGG (Kyoto Encyclopedia of Genes and Genomes [94]) pathway gene sets, limiting our search to metabolic and signaling functional annotations. Similarly, we obtained up-to-date gene ontology (GO) gene sets specifying biological processes. We mapped our FlyBase gene (FBgn) IDs to ENTREZ ids, and performed pathway and gene set enrichment, and GO analysis on the resulting gene list within GAGE with the `gage()` function. We tested for perturbation of each gene set relative to all genes in the experiment by calculating the mean individual statistics (`stat.mean`) from multiple gene set tests using a two-sample t-test implemented in GAGE as `gs.tTest()`, and obtain a global p-value from individual p-values. The global p-values were then adjusted for multiple testing using the Benjamini & Hochberg procedure [95], and refer to these as FDR [93].

#### Gene co-expression analysis

Next, we sought to identify sets of genes (modules) showing high correlation in their pattern of expression across tissues with respect to treatment condition. We used the `removeBatchEffect()` in the R limma package to correct for known (batch) and identified (SV) effects. We applied a regularized log transformation (rlog) to the batch-corrected matrix to minimize differences between samples for rows with low counts as well as normalize to library size. The rlog transformation is recommended if, as in this study (0.52 - 1.92), size factors vary widely [90]. The resulting expression matrix was used as input in the R package, Weighted Gene Co-expression Network Analysis, WGCNA, [96] to identify co-expressed gene modules showing similar patterns of expression across tissues and treatments. We built the initial network from samples over all treatments (N = 54) using a signed adjacency matrix with power 23 (i.e. function `pickSoftThreshold()`) to construct the topological overlap matrix (TOM) from all 12,614 genes (Fig. ADDITIONAL FILE 4). This power represents the lowest power for which the scale-free topology fit index curve flattens out after reaching a high value of r^2^ = 0.90. We performed hierarchical clustering of the TOM using the `flashClust()` function (method = “average”) which implements `hclust()` clustering more efficiently for large datasets [97]. We used the `cutTreeDynamic()` function to identify initial the initial set of modules at the following thresholds: `cutHeight = 0.95` (default), `deepSplit = 2, minClusterDendro = 30, pamRespectsDendro = FALSE`. A module eigengene is defined as the first principal component of a given set of co-expressed genes, and can be considered as a representative of the gene expression profiles in that set [96]. By convention, modules are referred to by their color labels, and, the grey module is used by default to contain genes not assigned to any module [98].

We then evaluated module membership using a resampling strategy described in Sikkink et al, [99]. Randomly, we chose four of six replicates for each diet-sample combination to produce 100 new datasets, each with 36 samples. Using the same parameters as in the full dataset, WGCNA was re-run for each new dataset and resulting modules examined for gene membership. First, a resampled module was accepted if it included at least 10% of the genes in the corresponding module in the full dataset. Secondly, a gene was considered to be strongly supported to belong to a module in the full dataset if it appeared in that module in at least 50% of resampled networks. We identified genes that failed to meet these criteria and placed them in the unassigned module. e merged highly correlated modules (*r* ⪰ 0.9) during network construction for each resampled dataset to be consistent with how the full dataset was analyzed.

To summarize major patterns of within-module expression with respect to diet condition, we extracted the first principal component of expression of genes in each module, called the module eigengene. We then performed an ANOVA for each module eigengenes to assess the effects of diet, tissue, and diet-tissue interactions on total module expression. Because all modules had a significant effect of either diet or diet-tissue interaction, we examined module enrichment for diet-related functional annotations. We therefore accessed Bioconductor annotation libraries `AnnotationDbi` and `org.Dm.eg.db` using the `GOenrichmentAnalysis()` function within WGCNA, and calculated the Fisher’s Exact test with the Bonferroni correction to identify significantly enriched GO terms in each module, providing all genes available in our experiment as a background list. We reviewed all terms significantly enriched (*P*_*adj.*_ < 0.01) for the BP category (to view terms for CC, MF, see Additional file 6). For this discussion, we restricted the analysis to the BP category to focus only on biological function and not biochemical activities (MF) or subcellular location where a gene product is active (CC), at the same time reducing the number of tests for enrichment as a way to limit the number of terms for interpretation.

## List of abbreviations

AMPK: Adenosine Monophosphate-activated Protein Kinase
ANOVA: Analysis of Variance
BP: Biological Process
C: Control
CC: Cellular Component
DR: Dietary Restriction
DSPR: *Drosophila* Synthetic Population Resources
eQTL: expression Quantitative Trait Locus
FDR: False Discovery Rate
GAGE: Generally Applicable Gene-set Enrichment for Pathway Analysis
GO: Gene Ontology
GSEA: Gene Set Enrichment Analysis
HS: High Sugar
IIS: insulin/insulin-like
KEGG: Kyoto Encyclopedia of Genes and Genomes
LRT: likelihood ratio test
MF: Molecular Function
mRNA: messenger Ribonucleic Acid
PCA: Principal Components Analysis
SVA: Surrogate Variable Analysis
TOM: Topological Overlap Matrix
TOR: target of rapamycin
WGCNA: Weighted Gene Co-expression Network Analysis

## Declarations

### Ethics approval and consent to participate

Not applicable

### Consent for publication

Not applicable

### Availability of data and materials

Trimmed read data are available in the Sequence Read Archive (SRA, SUB6076601) database (https://www.ncbi.nlm.nih.gov/sra) under the accession number XXX. Sample data, scripts and intermediate files are available here https://github.com/nochet/BasePop_RNAseq.

### Competing interests

The authors declare that they have no competing interests.

### Funding

Funding for this study was provided by NIH R01 GM117135 and NSF IOS 1654866 to EGK. Computational work was performed on the high-performance computing infrastructure provided by Research Computing Support Services and in part by the National Science Foundation under grant number CNS-1429294 at the University of Missouri, Columbia Mo.

### Authors’ contributions

EGK conceived and planned the project. EN and EGK designed, set up, and managed the experiment, AR isolated RNA and prepared samples for sequencing, EN, PAW-S and EGK analyzed the data, EN and EGK wrote the manuscript, EN, EGK, AR and PAW-S commented on the manuscript. All authors read and approved the final manuscript.

## Acknowledgments

We would like to acknowledge Elizabeth Lo Presti, Michael Reed, and Kevin Middleton for help with experimental setup and fly husbandry. High-throughput sequencing services were performed at the University of Missouri DNA Core Facility.

## Figures, table and additional files

Figure1_Study_Design.pdf

Figure2_Transcripts_summary_stats.pdf

Figure3_FoldChange_all.pdf

Figure4_VolcanoPlots_all.pdf

Figure5_Eigengenes_all.pdf

Figure6_Individualgenes.pdf

Figure7_QTL_FC.pdf

Additional_file_1_figures.docx

Additional_file_2_DEGs_lrt.treatment_0.05.csv

Additional_file_3_DEGs_lrt.interaction_0.05.csv

Additional_file_4_Table4_GSEA_sigPathways_sigGOTerms.xlsx

Additional file_5_Table5_WCGNA_eigengenes.csv

Additional_file_6_Table6_GOEnrich_WGCNA_Resamp.csv

Additional_file_7_Table7_DEGs_underQTL.csv

Additional_file_8_Table8_pathwayGenes_DE.csv

Additional_file_9_Table9_DietInfo.csv

Additional_file_10_Table10_SampleInfo_RNA.xlsx

Additional_file_11_Table11_SampleInfo_BatchInfo.xlsx

## Notes

#### Summary of Updates

This version is improved further for language, corrected for errors initially unseen, and includes referencing to further additional data files now provided. It is publication-ready.

