## Additional file 1 for "Diverse biological processes coordinate the transcriptional response to nutritional changes in a *Drosophila melanogaster* multiparent population"

Supplementary figures

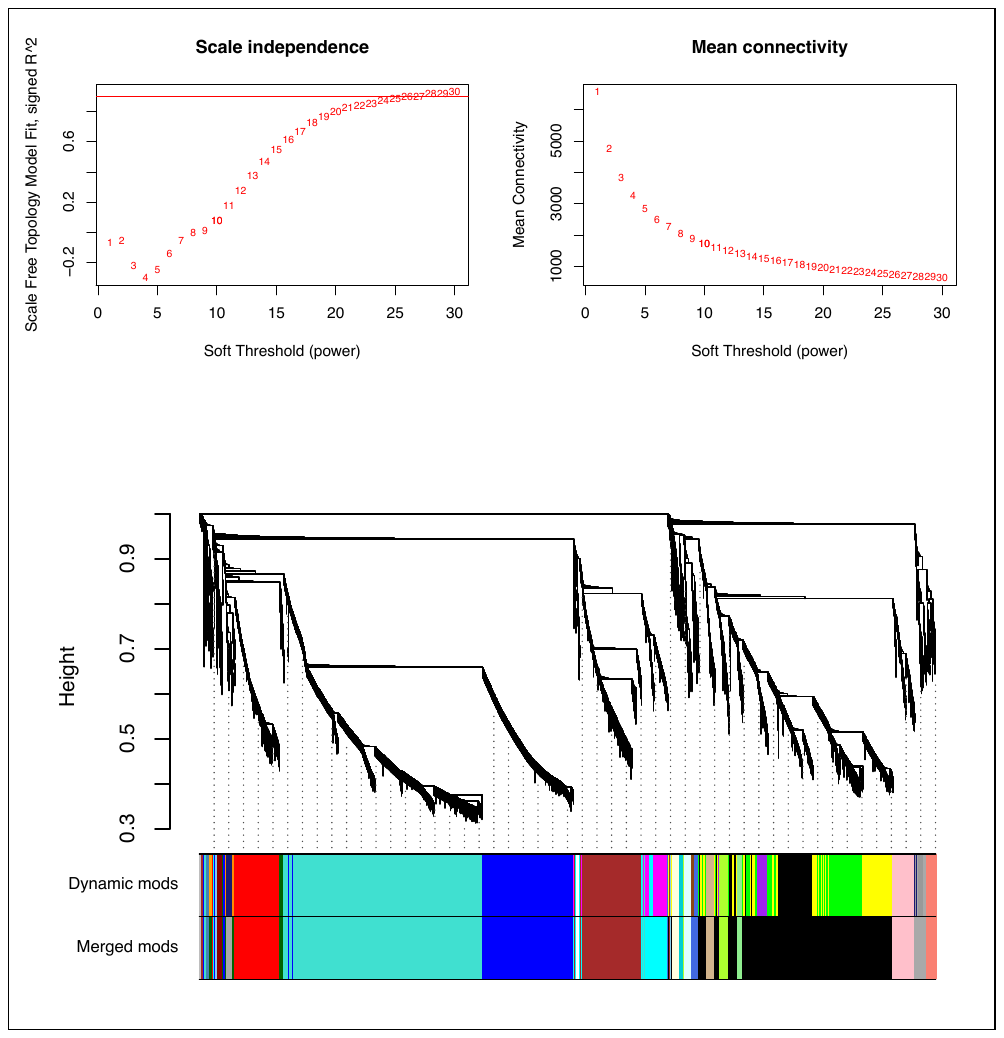

**Figure S1:** Module detection by hierarchical clustering of 12,614 genes showing co-expressed sets of gene clusters. *Top*: Selection of soft-threshold power – effect of soft-thresholding on the scale-free model fit. Because metabolic networks are likely to be scale-free, higher values of soft threshold power reduces noise of the correlations in the adjacency matrix. An optimal threshold value is one that maximizes network similarity to a scale-free graph - the lowest power for which the scale free topology index reaches 0.90 (i.e. 23 in this study, *top left*). *Top right* shows the effect of soft-threshold power on the mean connectivity. *Bottom*: Hierarchical clustering of genes using dissimilarity based on topological overlap. Height (*y*-axis*)* represents distance determined by the extent of topological overlap. A dendrogram depicts gene clusters (modules) detected. Each module is assigned a color name. Top color panel displays modules automatically detected with flashClust(). The lower panel depicts modules after merging highly correlated modules (*r* ⪰ 0.9).

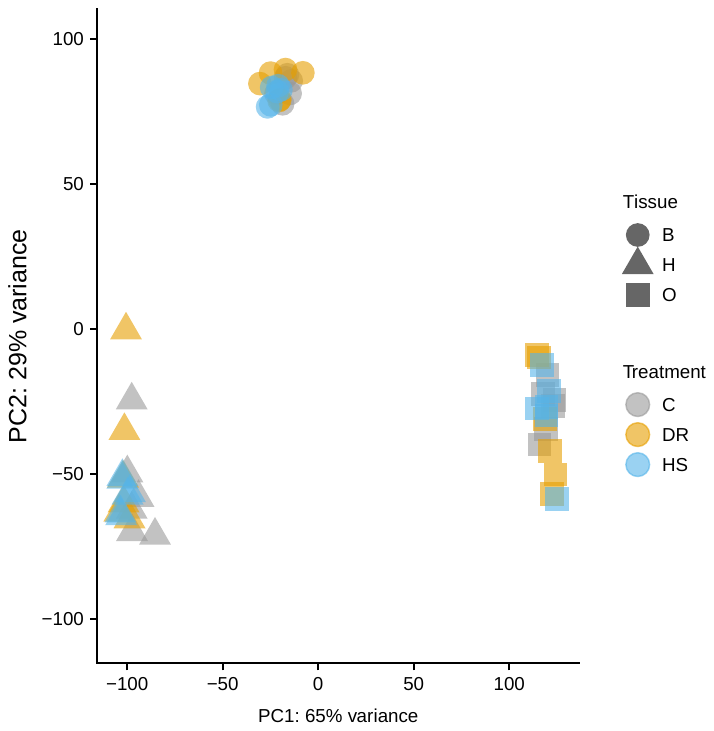

**Figure S2:** Principal components analysis (PCA) to visualize the overall effect of diet and tissue before correcting for batch effects (compare with Figure 2a in main text after batch effect correction). Two dimensions are shown (PC1 and PC2) accounting for 94% of the variance in sample gene expression.

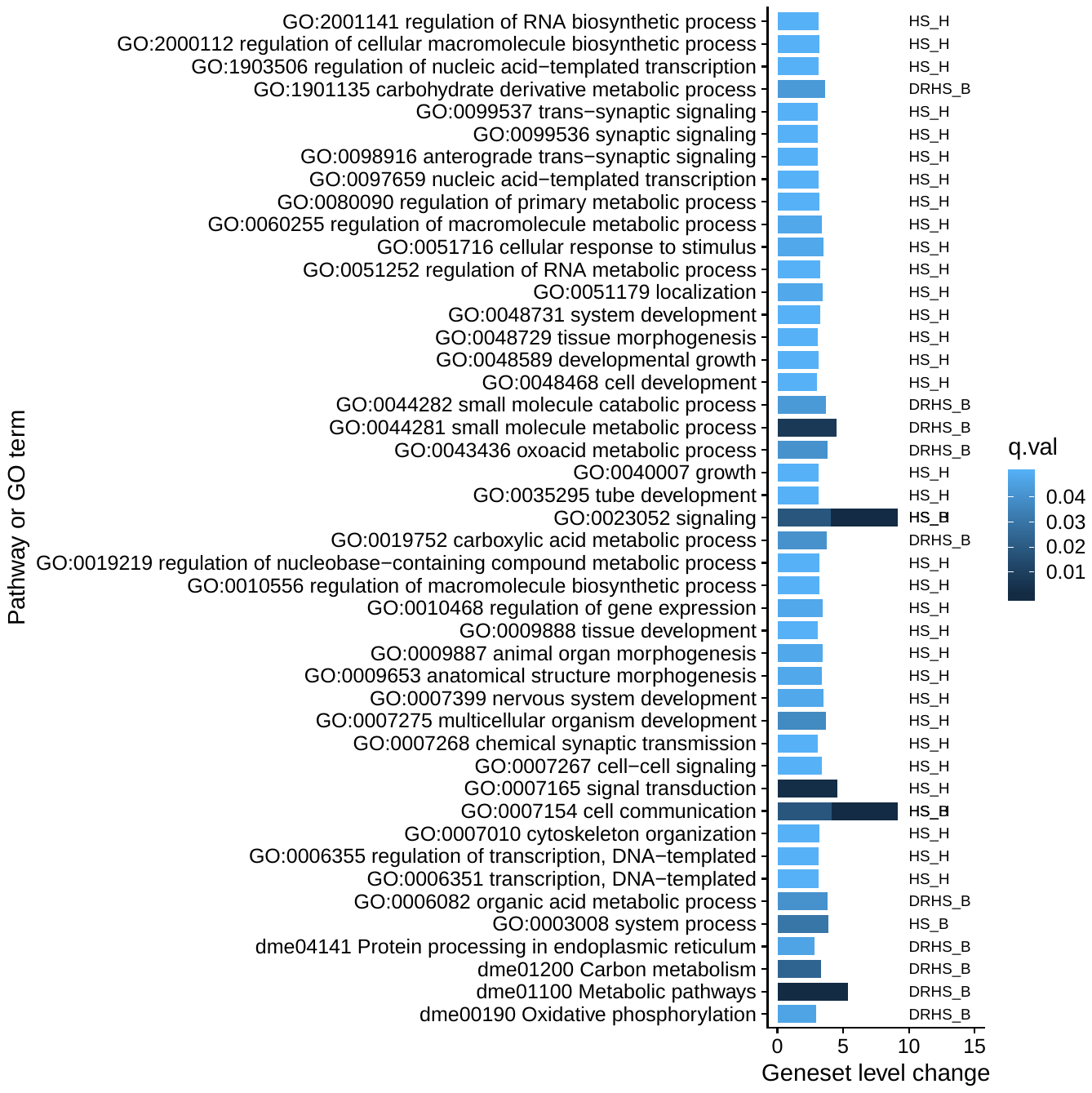

**Figure S3:** Gene set enrichment analysis (GSEA). Four pathways (starting with “dme”) and 41 gene ontology terms (GO) were identified from GSEA of the whole list significantly differentially expressed genes (2,475) for the main effect of diet. GSEA differs from traditional GO analysis in that genes are weighted by their fold change in the analysis of enrichment against pre-computed gene sets. This analysis identifies four pathways that are not any of the canonical pathways for the response to diet in model organisms such as IIS/TOR pathway. Secondly, a large number of GO terms represent many other biological processes (BP) in addition to nutrient metabolism. Clearly, in this population, the response to diet is not limited to canonical nutrient sensing pathways.
