## Additional file 9 for "Diverse biological processes coordinate the transcriptional response to nutritional changes in a *Drosophila melanogaster* multiparent population"

Table S9: Composition of the four diets used in this experiment.

|  | **Maintenance** | **DR** | **C** | **HS** |
| --- | --- | --- | --- | --- |
| **Water (ml)** | 1066 | 1000 | 1000 | 1000 |
| **Agar (g)** | 6.25 | 10 | 10 | 10 |
| **Dextrose (g)** | 86.26 | - | - | - |
| **Sucrose (g)** | - | 50 | 50 | 342 |
| ***Molarity*** | *-* | 0.15 | 0.15 | 1 |
| **Yeast (g)** | 21.6 | 100 | 200 | 200 |
| **Cornmeal (g)** | 40.8 | - | - | - |
| **Tegosept (g)** | 1.8 | 2.7 | 2.7 | 2.7 |
| **Ethanol (ml)** | 7.3 | 11 | 11 | 11 |
| **% Protein** | 10-13 | 36-41 | 45-53 | 17-19 |
| **% Carbohydrate** | 93-95 | 59-64 | 47-52 | 81-83 |
